# Genomic diversity of SARS-CoV-2 can be accelerated by mutations in the nsp14 gene

**DOI:** 10.1101/2020.12.23.424231

**Authors:** Kosuke Takada, Mahoko Takahashi Ueda, Shintaro Shichinohe, Yurie Kida, Chikako Ono, Yoshiharu Matsuura, Tokiko Watanabe, So Nakagawa

## Abstract

Coronaviruses, including severe acute respiratory syndrome coronavirus 2 (SARS-CoV-2), encode a proofreading exonuclease, nonstructural protein 14 (nsp14), that helps ensure replication competence at a low evolutionary rate compared with other RNA viruses. In the current pandemic, SARS-CoV-2 has accumulated diverse genomic mutations including in nsp14. Here, to clarify whether amino acid substitutions in nsp14 affect the genomic diversity and evolution of SARS-CoV-2, we searched for amino acid substitutions in nature that may interfere with nsp14 function. We found that viruses carrying a proline-to-leucine change at position 203 (P203L) have a high evolutionary rate and that a recombinant SARS-CoV-2 virus with the P203L mutation acquired more diverse genomic mutations than wild-type virus during its replication in hamsters. Our findings suggest that substitutions, such as P203L, in nsp14 may accelerate the genomic diversity of SARS-CoV-2, contributing to virus evolution during the pandemic.

## Introduction

Virus mutates rapidly, resulting in the increased genetic diversity that drives virus evolution and facilitates virus adaptation. Generally, RNA viruses have a high mutation rate because they replicate their genome by using an error-prone RNA-dependent RNA polymerase (RdRp) so that nucleotide incorporation errors are not corrected (1-3). In contrast, coronaviruses have lower mutation rates because they possess nonstructural protein 14 (nsp14), which contains an exoribonuclease (ExoN) domain that provide a proofreading function (4, 5). Nonetheless, in the case of severe acute respiratory syndrome coronavirus-2 (SARS-CoV-2), which emerged in China at the end of 2019 and caused the COVID-19 pandemic, numerous variants have emerged by acquiring multiple mutations in a short period of time. Some of these variants have been a cause for concern due to their increased infectivity, transmissibility, pathogenicity, and/or reduced sensitivity to vaccines and therapeutic agents. These variants, classified as ‘variants of concern (VOCs)’ by the World Health Organization (WHO), include Alpha (PANGO ID B.1.1.7), Beta (PANGO ID B.1.351), Gamma (PANGO ID P.1), Delta (PANGO ID B.1.617.2), and most recently Omicron (PANGO ID B.1.1.529). To control COVID-19 and bring the pandemic to an end, it is important to understand the mechanism of the emergence of these diverse variants and the genomic evolution of SARS-CoV-2.

Coronaviruses possess a positive-sense, single-stranded RNA genome that is one of the largest among known RNA viruses, ranging from 26 to 32 kb (2). There are two large open reading frames (ORFs) at the 5′ proximal end, ORF1a and ORF1b, which comprise approximately two-thirds of the genome, and encode two large polyproteins, pp1a and pp1ab, that are cleaved into about 16 nonstructural proteins (nsps). Some of the processed nsps assemble to form the replication-transcription complex, which contains an RdRp (nsp12), the helicase (nsp13), processivity factors (nsp7 and nsp8), and the proofreading ExoN complex (nsp14 and nsp10) (6-8). During genome replication, nsp14 ExoN activity plays a role in the RNA proofreading machinery by efficiently removing mismatched 3’-endnucleotides (9). Experiments in which the nsp14 of SARS-CoV was functionally inactivated revealed 10–20 times higher mutation rates (5, 10, 11), indicating that nsp14 may be essential for maintaining the virus genome.

In this study, to clarify whether amino acid substitutions in nsp14 affect the genomic diversity and evolution of SARS-CoV-2, we searched for amino acid substitutions in nature that may interfere with the function of nsp14 in SARS-CoV-2. First, to obtain functionally important sites in nsp14, we examined 62 representative coronaviruses belonging to the family *Coronaviridae* and found that 99 of the 527 amino acid sites of nsp14 were evolutionarily conserved. We then examined nsp14 sequences obtained from 28,082 SARS-CoV-2 genomes available in the GISAID EpiCoV database, which includes not only sequencing data but also sampling date, country, and institution information (12), and identified an additional 6 amino acid changes in nsp14 mutants that were not detected in the 62 representative coronaviruses. We examined the genome mutation rates of these mutants and found that the amino acid replacement of proline (P) for leucine (L) at position 203 of nsp14 could be involved in increasing the nucleotide mutation rate of SARS-CoV-2 genomes. To investigate the biological significance of the nsp14 P203L substitution, we generated recombinant SARS-CoV-2 viruses possessing wild-type nsp14 or mutant nsp14 with P203L and characterized them in a hamster model. We found that the nsp14-P203L mutant grown in the lungs of hamsters acquired significantly more diverse genomic mutations than did the wild-type virus. These results indicate that substitutions in nsp14, such as P203L, could accelerate the genomic diversity of SARS-CoV-2.

## Results

### Conserved regions of nsp14 proteins among coronaviruses

We first generated a phylogenetic tree of the amino acid sequences of the nsp14 gene obtained from 62 representative coronaviruses (the viruses used in this analysis are summarized in Supplementary Table 1). The phylogenetic tree clearly showed the relationship among the four genera of coronaviruses (Fig. 1A), consistent with results obtained by using the partial amino acid sequences of ORF1ab described in a previous report (13). Our results suggest that the nsp14 gene is broadly conserved among coronaviruses, and that the conserved amino acid sites could be important for its molecular function.

**Fig. 1:**
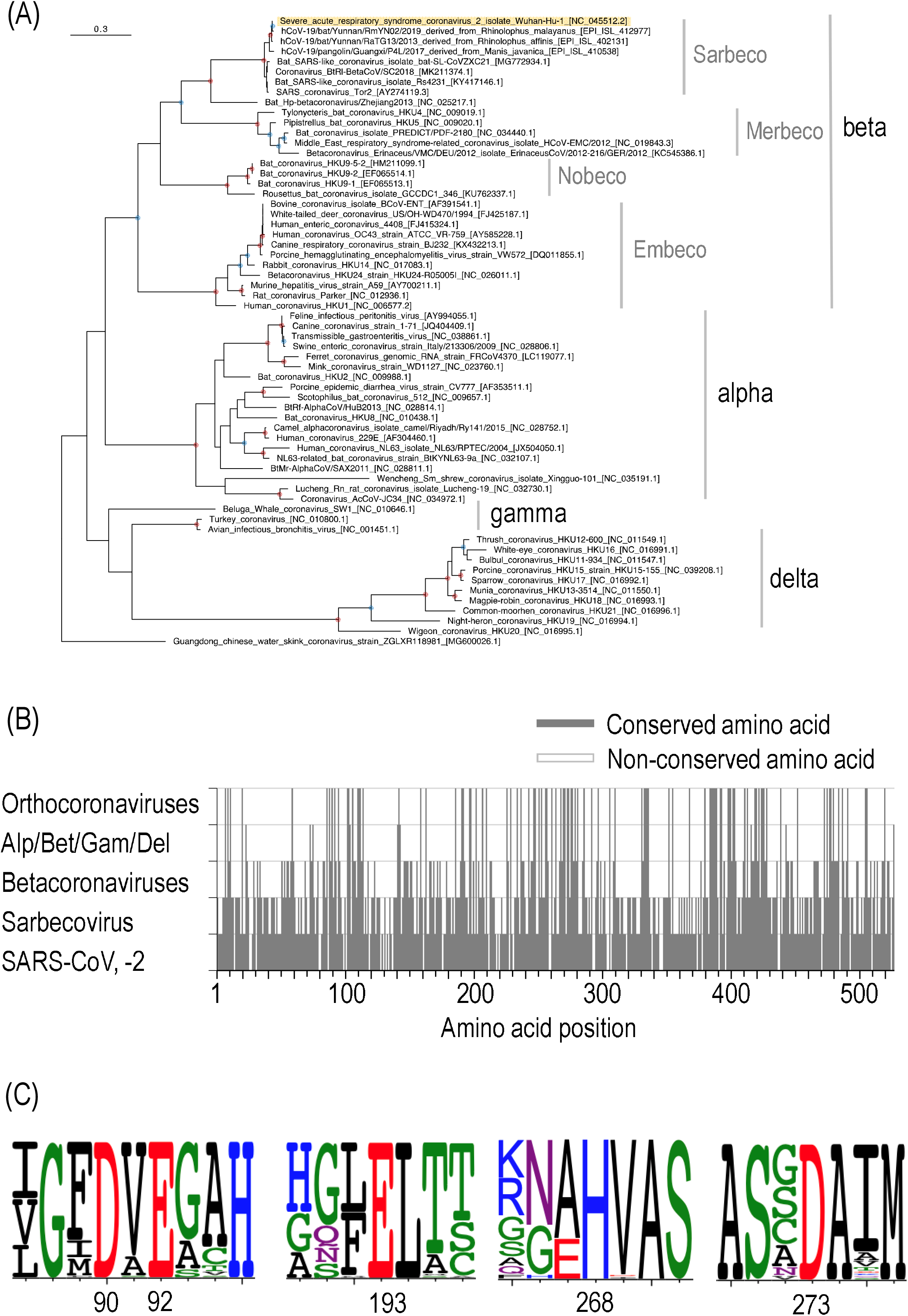
Phylogenetic tree and amino acid sequence alignment of nsp14 derived from representative coronaviruses. (A) Maximum likelihood (ML)-based phylogenetic tree of the 62 representative coronaviruses. Amino acid sequences of nsp14 in ORF1ab were used for this analysis. A red or blue circle in an internal node corresponds to bootstrap values ≥ 95% or ≥ 80 %, respectively. The virus located at the top is SARS-CoV-2 (hCoV-19/Wuhan/Hu-1/2019). (B) Conserved amino acid residues among each group. The amino acid sequence of the nsp14 of the 62 representative coronaviruses is shown according to the relative amino acid positions of SARS-CoV-2 nsp14 (note that the gaps in the SARS-CoV-2 sequence are not shown). The conserved amino acid positions are indicated in gray. (C) Five conserved active site residues involved in proofreading. The amino acid proportions for each site were calculated based on the sequence alignment of the nsp14 of the 62 representative coronaviruses visualized by using WebLogo (27). The amino acids are colored according to their chemical properties.

Next, we examined the conserved amino acid sites of the nsp14 gene among the 62 coronaviruses (Fig. 1B). The amino acid positions of nsp14 were based on the SARS-CoV three-dimensional structure (PDB ID: 5C8S); SARS-CoV and SARS-CoV-2 both belong to the subgenus *Sarbecovirus* in the genus *Betacoronavirus*. For nsp14, the number of identical amino acids was 482/527 (91.5%) in 8 representative coronaviruses belonging to the subgenus *Sarbecovirus*, and 359/527 (68.1%) if we included the nsp14 of the Bat Hp-betacoronavirus/Zhejiang2013 (GenBank ID: NC_025217.1) belonging to the subgenus *Hibecovirus*, an outgroup of *Sarbecovirus*. Moreover, 200/527 (38.0%) positions were conserved among 29 viruses of the genus *Betacoronavirus*, and 108/527 (20.5%) positions were conserved among the 61 coronaviruses (Fig. 1B). If we included an unclassified coronavirus found in *Tropidophorus sinicus* (Chinese waterside skink) (14), the conserved amino acid positions were 99/527 (18.8%). These results suggest that these amino acid sites of the nsp14 gene are conserved among diverse coronaviruses because they may be functionally important. We further validated this hypothesis by calculating nonsynonymous (dN) and synonymous (dS) substitution rates on a per-site basis for the nsp14 sequences of the 62 coronaviruses by using the SLAC program (see Materials and Methods) (15). For the dN/dS analysis, we removed 23 amino acid sites that contained insertions and deletions out of the total 527 amino acid sites in the SARS-CoV-2 nsp14 of 61 coronaviruses (excluding the Chinese waterside skink coronavirus that contains a mixture of nucleotides at these positions). Thus, 504 amino acid sites of the 61 coronavirus nsp14 multiple alignment were calculated for the dN/dS ratios; no positive selection sites were found in nsp14, whereas 379 of the 504 codon sites were under negative selection (*P* < 0.01, Supplementary Fig. 1). In total, 91 amino acids were unchanged and under negative selection in the nsp14 of the 62 representative coronaviruses.

We then investigated whether the amino acids that are known to be important for the error-correcting function of nsp14 are conserved. Coronavirus nsp14 possesses both ExoN and N7-MTase activities. The ExoN domain of coronavirus nsp14 was originally identified based on its sequence similarity to distant cell homologs (9) and has been assigned to the DEDD exonuclease superfamily, which includes the proofreading domains of many DNA polymerases and other eukaryotic and prokaryotic exonucleases. The nsp14 proteins have four conserved active site residues distributed in three standard motifs of the primary structure: residues 90D/92E (motif I), 191E (motif II), and 268H/273D (motif III) (16-18). We found that the five active sites of nsp14 were conserved among the 62 coronaviruses examined in this study (Fig. 1C). Moreover, there were three zinc fingers that were essential for nsp14 function: zinc finger 1 (ZF1) comprising 207C, 210C, 226C, and 229H; zinc finger 2 (ZF2) comprising 257H, 261C, 264H, and 279C; and zinc finger 3 (ZF3) comprising 452C, 473C, 484C, and 487H. S-adenosyl methionine (SAM)-binding motif I includes residues 331D, 333G, 335P, and 337A (or 337G), which are involved in the (N7 guanine)-methyl transferase reaction by nsp14. We found that all these residues were conserved among the nsp14 sequences of the 62 coronaviruses examined (Supplementary Fig. 2). These results suggest that the amino acids of nsp14 that are functionally important are conserved among the diversified coronaviruses.

### Diversity of nsp14 in SARS-CoV-2

Amino acid comparisons of nsp14 in the 62 representative coronaviruses revealed that several sites are strongly conserved and that some substitutions may have deleterious effects on the survival of SARS-CoV-2. SARS-CoV-2 was first detected in humans in 2019 and has spread rapidly all over the world in a short period of time. Therefore, some nsp14 mutations that impair its error-correcting function may remain in the SARS-CoV-2 population. Accordingly, we attempted to identify such deleterious mutations in the current SARS-CoV-2 population.

We downloaded the aligned nucleotide sequences of SARS-CoV-2 (87,625 sequences) from the GISAID database as of September 7, 2020. The SARS-CoV-2 genomes isolated from humans were extracted (87,304 sequences), and the sequences containing undetermined and/or mixed nucleotides were removed; the remaining 28,082 sequences were used in subsequent analyses. The coding region of nsp14 was translated and compared. The results showed that 25,913 nsp14 amino acid sequences (92.27%) were identical to that of the reference sequence, whereas 2169 sequences contained at least one amino acid substitution in nsp14 (Supplementary Table 2). The top 20 variants possessing nsp14 with amino acids that were different from the major sequence (shown in Supplementary Fig. 2) are summarized in Table 1. Note that frameshift mutants were excluded from the table because they may affect downstream genes as well. Notably, among the 20 amino acid nsp14 mutants, six amino acids (177F, 377L, 369F, 501I, 203L and 297S) were not found in the 62 nsp14 proteins of the representative coronaviruses (Supplementary Fig. 2). We further analyzed the evolutionary selection pressures of these 6 amino acid sites by using 73 coronaviruses belonging to the genus *Betacoronaviruses* that includes SARS-CoV-2 (Supplementary Table 3). We confirmed that the six amino acid variants were not found in any of the 73 Beta-coronaviruses (Supplementary Table 3). The six conserved positions (177, 203, 297, 369, 377, and 501) were also found to be under negative selection (*P* < 0.01, Supplementary Table 4). Overall, these results indicate that the conserved amino acid sites of nsp14 are also evolutionarily selected in the genus *Betacoronaviruses*.

### Correlation between the amino acid sequences of nsp14 and nucleotide mutation rates in the viral genome

To examine whether any of the six amino acid substitutions in the nsp14 of SARS-CoV-2 variants could affect its proofreading function, we looked for a correlation between the amino acid variants of nsp14 and the genetic diversities of SARS-CoV-2 genomes. For this purpose, we roughly estimated evolutionary rates of SARS-CoV-2 possessing mutated nsp14, compared to that of SARS-CoV-2 with wild-type nsp14. Sequences for which sampling date information (year, month, and day) was not available were excluded from this analysis. Nucleotide mutations per year were calculated based on the regression coefficient using hCoV-19/Wuhan/Hu-1/2019 (GISAID ID: EPI_ISL_402125) as the reference sequence (see Materials and Methods). Note that the estimated evolutionary rates of each nsp14 mutant were not phylogenetically corrected; the estimated rates were used only to obtain nsp14 mutation candidate(s) that may affect evolutionary rates.

We found that SARS-CoV-2 containing wild-type nsp14 had 19.8 nucleotide mutations in its whole genome per year (Table 2; Supplementary Fig. 3 for the regression coefficient). Conversely, seven SARS-CoV-2 variants containing nsp14 with amino acid substitutions showed 27.3 (L177F), 59.8 [L177F/T372I (*i*.*e*., L177F and T372I)], 35.9 (P203L), 17.1 (P297S), 26.2 (S369F), 13.0 (F377L), or 21.9 (M501I) nucleotide mutations per year (Table 2; Supplementary Fig. 3). Although the substitution rates of the L177F/T372I and F377L variants were found to be unreliable because of limited time ranges, those of the other variants were correlated (Supplementary Fig. 3). Therefore, SARS-CoV-2 variants possessing nsp14 with the L177F, P203L, S369F, or M501I substitution may have a higher genomic mutation rate than that of SARS-CoV-2 carrying wild-type nsp14. In particular, the P203L variant of nsp14 (nsp14-P203L), which showed the highest mutation rate (35.7 nucleotide mutations per year), is a strong candidate that enhances the diversity of SARS-CoV-2 genomes.

**Table 1.** Amino acid sequence of nsp14 in SARS-CoV-2 epidemic strains

| Appearance frequency ranking | Substitution in nsp14 | Number of sequences | Frequency of appearance (%) |
| --- | --- | --- | --- |
| 1 | — | 25913 | 92.27 |
| 2 | A320V | 192 | 0.68 |
| 3 | F233L | 144 | 0.51 |
| 4 | S374A | 123 | 0.44 |
| **5** | **L177F** | **116** | **0.41** |
| 6 | T250I | 61 | 0.22 |
| **7** | **F377L** | **51** | **0.18** |
| 8 | A138V | 39 | 0.14 |
| 9 | V120A | 35 | 0.12 |
| 10 | T16I | 34 | 0.12 |
| **11** | **S369F** | **33** | **0.12** |
| 12 | G481S | 31 | 0.11 |
| **13** | **M501I** | **28** | **0.10** |
| 13 | D345G | 28 | 0.10 |
| 13 | N129D | 28 | 0.10 |
| **16** | **P203L** | **26** | **0.09** |
| **16** | **L177F/T372I** | **26** | **0.09** |
| **18** | **P297S** | **25** | **0.09** |
| 18 | T31I | 25 | 0.09 |
| 20 | D324Y | 24 | 0.09 |

28,082 SARS-CoV-2 genome sequences that were annotated as Human and free of undetermined or mixed nucleotides were used in the analysis. Amino acid substitutions were defined as a change relative to the hCoV-19/Wuhan/Hu-1/2019 reference sequence. Boldface characters indicate that the amino acid replacement was not observed in the 62 coronaviruses examined in this analysis. —, no mutation was found compared with the reference sequence.

**Table 2.** Nucleotide mutation rates of SARS-CoV-2 with different nsp14 variants

| Ranking of nsp14 | Substitution in nsp14 | Number of<br>comparison sequences | Substitution rate per year in the whole<br>genome |
| --- | --- | --- | --- |
| 1 | — | 25471 | 19.8 |
| 5 | L177F | 114 | 27.3 |
| 7 | F377L | 51 | 13.0 |
| 11 | S369F | 33 | 26.2 |
| 13 | M501I | 28 | 21.9 |
| 16 | P203L | 24 | 35.9 |
| 17 | L177F/T372I | 26 | 59.8 |
| 18 | P297S | 25 | 17.1 |

The number of mismatched nucleotides in the alignment of the reference sequence with the comparison sequence of each group was counted, and the average value was calculated. hCoV-19/Wuhan/Hu-1/2019 was selected as the reference sequence. Note that the estimated substitution rates were not phylogenetically corrected. —, no mutation was found compared with the hCoV-19/Wuhan/Hu-1/2019 sequence.

We then verified the substitution rates of the nsp14-P203L variants for each single cluster in which the maximum number was observed: 6 genomes for the nsp14-203L mutant and 194 genomes for the nsp14-203P wildtype in the B.1.1.33 lineage (Fig. 2A; Supplementary Fig. 4 with GISAID IDs). We found that nsp14-P203L variants in the B.1.1.33 lineage forming a single clade showed a high nucleotide evolutionary rate (58.8 nucleotide mutations/year) compared with those with the nsp14-203P (*i*.*e*., wildtype) strains in the same lineage (16.5 nucleotide mutations/year) (Fig. 2B and Table 3). Although the number of variants was limited (6), a correlation was clearly observed (Fig. 2B). In addition, six other nsp-P203L variants in a different cluster (B.1) also showed a relatively high nucleotide mutation rate (28.0 nucleotide mutations/year), assuming that variants were in the same clade if their PANGO IDs (19) and GISAID clades were identical (Table 3; Supplementary Fig. 5 for regression coefficients). With the same assumption, we did the same calculation for the other nsp14 mutants in the lineages (Table 3; Supplementary Fig. 5) and found that nsp14-P203L in the B.1.1.33 lineage exhibited the highest substitution rate. These results indicate that SARS-CoV-2 variants containing nsp14-P203L are strong candidates for variants that readily accumulate mutations in the viral population.

**Table 3.** Nucleotide mutation rates of the seven nsp14 variants for each cluster in which the maximum number was observed

| Ranking<br>of nsp14 | Substitution in<br>nsp14 | Lineage (PANGO ID) | Number of sequences<br>containing the nsp14<br>substitution | Substitution rate per year in<br>the whole genome |
| --- | --- | --- | --- | --- |
| 5 | L177F | B.1.1.25 | 54 | 7.8 |
| 7 | F377L | B | 50 | 8.5 |
| 11 | S369F | B.1.1 | 17 | 13.4 |
| 13 | M501I | B.1 | 11 | 4.4 |
| 16 | P203L | B.1.1.33 | 6 | 58.8 |
|  |  | B.1 | 6 | 28.0 |
| 17 | L177F/T372I | B.1.113 | 20 | 50.1 |
| 18 | P297S | B.4 | 11 | 16.1 |

**Fig. 2:**
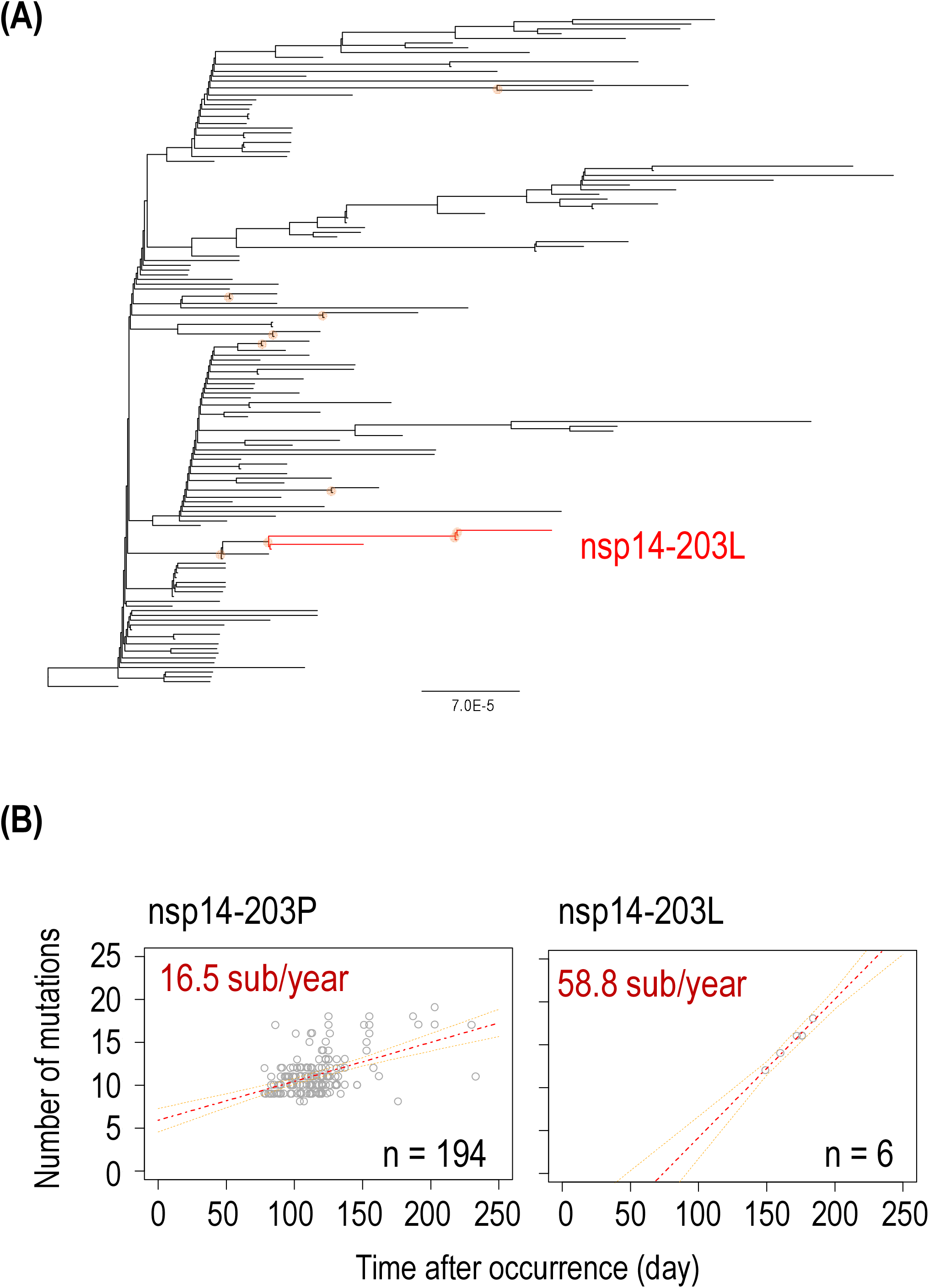
Phylogeny and nucleotide mutation rate of nsp14-P203L variants in the PANGO lineage B.1.1.33. (A) ML tree of SARS-CoV-2 genomes in the B.1.1.33-lineage. The nsp14-P203L variant is highlighted in red. An orange circle corresponds to bootstrap values ≥ 70%. (B) Nucleotide mutation rates of nsp14-203P or nsp14-203L variants belonging to the B.1.1.33-lineage. All sample dots are displayed. Mutation rates per year in the genome are shown in red letters. The red and orange dotted lines correspond to the regression line and 95% confidence intervals, respectively.

The frequency of nucleotide mutations of coronaviruses may be affected not only by the nsp14 proofreading mechanism but also by the characteristics of other proteins, such as the fidelity of the RNA-dependent RNA polymerase (nonstructural protein 12; nsp12) (20). It is possible that amino acid substitutions in other proteins involved in genome replication affect the nucleotide mutation rates of SARS-CoV-2. Therefore, we examined amino acid substitutions among nsp14-203L variants involving the following genes that participate in gene replication: nsp7, nsp8, nsp9, nsp10, nsp12, nsp13, nsp15, and nsp16. The results showed that a proline-to-leucine substitution at position 323 (P323L) of nsp12 frequently occurred (20/25) in sequences of SARS-CoV-2 possessing nsp14-203L (Supplementary Table 5). However, the P323L mutation in nsp12 (nsp12-P323L) is known to accompany an aspartate-to-glycine substitution at position 614 (D614G) of S protein (21, 22). The S-D614G mutant was first reported at the end of January 2020 in China and Germany, and by March 2020, this mutant was a major variant in various regions all over the world (21). The P323L mutation in nsp12 was found in nsp14-203L variants as well. Indeed, the genomic mutation rate of SARS-CoV-2 with the P323L amino acid substitution in nsp12, was found to be 17.7 nucleotide mutations per year (Supplementary Fig. 6). Therefore, it is unlikely that P323L in nsp12 directly affects the genomic mutation rate of SARS-CoV-2 viruses carrying the P203L amino acid substitution in nsp14.

### The nsp14-P203L variant in the SARS-CoV-2 population

The nsp14-P203L variants have been isolated from 26 patients mainly in Europe and North America, with the first being reported on March 5, 2020 in The Netherlands (GISAID ID: EPI_ISL_454756, Supplementary Table 6). Considering their PANGO lineages (19) and GISAID clades (shown in Supplementary Table 6), the nsp14-P203L variants appeared at least 10 times. In fact, the nsp14-P203L substitution was observed in both Spike D614 strains and G614 variants, which have enhanced infection efficiency and have spread worldwide (21). However, we also found that the prevalence of nsp14-P203L variants did not increase steadily over time; the largest number of nsp14-P203L variants found in the B.1.1.33 and B.1 lineages was only six for each (Supplementary Table 6).

### The nsp14-P203L mutation effect on protein structure

To assess the effect of the P203L mutation of nps14, we first investigated the location of residue 203 by using a tertiary structure model of SARS-CoV-2 nsp14 (PDB ID: 7EGQ) (23). The residue 203 is exposed on the surface of the protein and is in the loop stretching from residues 200 to 214, which is in the interstices of the zinc and magnesium binding sites (positions 207 and 191). We found that the loop is located at the edge of an interaction surface for binding to the co-factor SARS-CoV-2 nsp10 (Fig. 3B and Supplementary Fig. 7A). The interaction with nsp10 has been shown to greatly increase the nucleolytic activity of SARS-CoV nsp14 (24). Therefore, to investigate the possibility that residues at or around position 203 play an important role in the nsp10-nsp14 interaction, we calculated the change in binding free energy (ΔΔGbind) induced by nsp14-P203L by using the BeQtMuSicC program. The predicted result showed that this substitution decreases binding affinity (ΔΔGbind 0.43 kcal/mol, Supplementary Fig. 7B), suggesting that the residue at position 203 is important to establish the interaction interface for binding to nsp10.

**Fig. 3:**
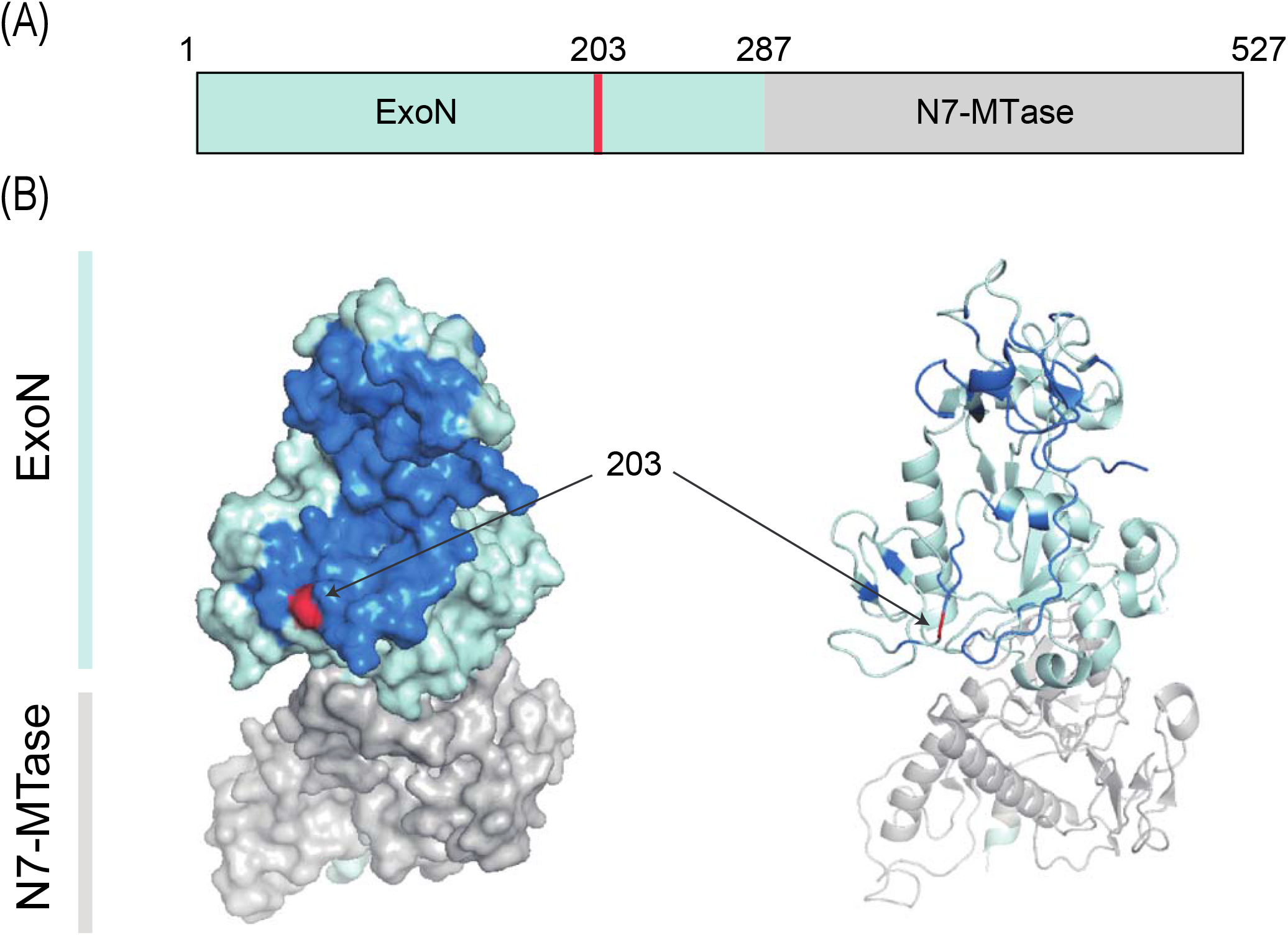
Three-dimensional position of the amino acid at position 203 in nsp14. (A) Domain structure of nsp14 with residue numbers. The ExoN and N7-Mtase domains are shown in light blue and gray, respectively. The amino acid at position 203 in nsp14 is shown in magenta. (B) Schematic representation of nsp14. The coloring is the same as in (A). The site of interaction with nsp10 is shown in dark blue.

### Characterization of a recombinant SARS-CoV-2 virus possessing nsp14-P203L *in vitro*

Genome data analysis suggested that SARS-CoV-2 variants containing nsp14-P203L could accumulate mutations in the viral population. Does the nsp14-P203L amino acid substitution affect viral properties (e.g., increasing the diversity of the viral genome)? To attempt to answer this question, we generated recombinant SARS-CoV-2 (based on hCoV-19/Japan/TY-WK-521/2020) viruses possessing wild-type nsp14 or P203L nsp14 by using a recently established circular polymerase extension reaction (CPER) method (25). We then we examined the effect of the nsp14-P203L amino acid substitution on the biological properties of SARS-CoV-2.

To determine whether the nsp14-P203L amino acid substitution affects the viral growth efficiency in cultured cells, we examined viral replication efficiency in VeroE6/TMPRSS2 cells. We found that both wild-type virus and nsp14-P203L viruses grew well, with max titers (mean titers = 7.35 log unit and 7.01 log unit, respectively) at 36 h post-infection (Fig. 4A), although the titer of the wild-type virus at 24 h post-infection was slightly higher than that of the nsp14-P203L virus (Fig. 4A). Next, we analyzed the genomic diversity of the recombinant virus grown in the cultured cells.

**Fig. 4:**
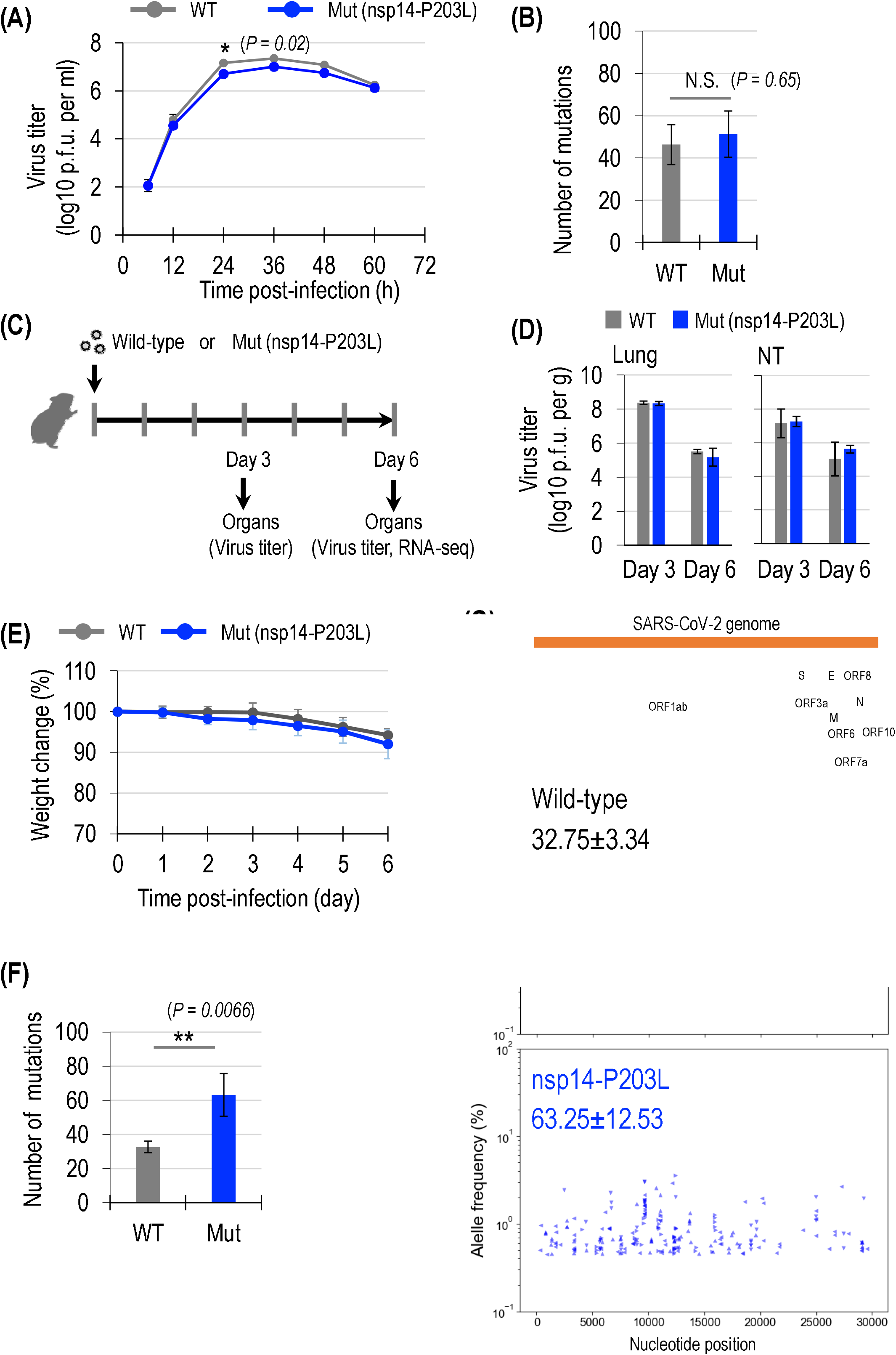
Characterization of the recombinant nsp14-P203L virus *in vitro* and *in vivo*. (A and B) Growth efficiency and genomic mutations of nsp14-P203L variants in cultured cells. (A) Growth kinetics of recombinant viruses in VeroE6/TMPRSS2 cells. VeroE6/TMPRSS2 cells were infected with viruses at a multiplicity of infection of 0.001. The supernatants of the infected cells were harvested at the indicated times, and virus titers were determined by the using plaque assays in VeroE6/TMPRSS2 cells. Data are shown as the mean of three independent experiments done in triplicate. Error bars indicate the standard deviation. Mean values were compared by using a two-way ANOVA, followed by Dunnett’s test (\**P*<0.05). Asterisks indicate statistically significant differences between WT and Mut viruses. (B) Number of genomic mutation types detected in viruses grown in VeroE6/TMPRSS2 cells. Three wild-type or mutant viruses were each plaque-purified, and each viral clone was infected into VeroE6/TMPRSS2 cells; supernatants were collected after 48 hours. RNA was extracted from the virus in the supernatant and RNA sequencing was performed. Mutations with a read depth of at least 1000x and an allele frequency of at least 0.5% were compared between the wild-type and mutant viruses. Mean values were compared by using an unpaired Student’s t test (N.D.; *P*>0.05). (C, D, E, F and G) Growth efficiency and genomic mutations of nsp14-P203L variants in Syrian hamsters. (C) Schematic overview of the animal experiment. Syrian hamsters were inoculated with 103 PFU of wild-type or nsp14-P203L virus via the intranasal (30 μL) route. (D) Virus replication in infected Syrian hamsters. Four Syrian hamsters per group were killed on days 3 and 6 post-infection for virus titration. Virus titers in lung or nasal turbinate (NT) organs were determined by using plaque assays in VeroE6/TMPRSS2 cells. Error bars indicate the standard deviation. Mean values were compared by using a two-way ANOVA, followed by Dunnett’s test. (E) Body weight changes in Syrian hamsters after viral infection. Body weights of virus-infected (n = 8) animals were monitored daily for 3 days. Body weights of virus-infected (n = 4) animals were monitored daily for 6 days. Data are presented as the mean percentages of the starting weight (±SD). (F) Number of genomic mutation types detected in viruses grown in Syrian hamster lung. RNA was extracted from the virus in lung homogenates from the Day 6 post-infection Syrian hamsters, and RNA sequencing was performed. Mutations with a read depth of at least 1000x and an allele frequency of at least 0.5% were compared between the wild-type and mutant viruses. Mean values compared by using an unpaired Student’s t test (**P<0.01). (G) SARS-CoV-2 genome organization and genomic locations of synonymous or nonsynonymous substitutions detected in the genomes of viruses grown in hamster lungs. The different directions of the triangles indicate each of the four individual hamsters. The number of nucleotide changes in viruses grown in Syrian hamster lung. The number of nucleotide changes, detected as synonymous and nonsynonymous substitutions, at a read depth of at least 1000x and an allele frequency of >0.5% were compared between the wild-type and nsp14-P203L mutant viruses.

Three plaque-purified wild-type or nsp14-P203L virus clones were used to infect VeroE6/TMPRSS2 cells, and the genomes of the viruses in the supernatant were collected 48 h later and analyzed by RNA sequencing. Mutations with a read depth of at least 1000 times and an allele frequency of at least 0.5% were compared between the wild-type and nsp14-P203L mutant viruses. The nsp14-P203L mutant grown in VeroE6/TMPRSS2 had 51.33 (±10.96) mutations and the wild-type virus had 46.33 (±9.46) mutations, which was not significantly different (*P =* 0.65) (Fig. 4B). These results suggest that nsp14-P203L allows efficient virus replication and may not affect viral genome mutagenesis in culture cells.

### Biological properties of the recombinant nsp14-P203L virus *in vivo*

Next, to examine whether the nsp14-P203L amino acid substitution affects viral properties *in vivo* (e.g., increasing viral replication efficiency and/or viral genome diversity), we investigated the biological properties of the nsp14-P203L mutant in a hamster model (Fig. 4C). We inoculated the hamsters intranasally with 10^3^ PFU of wild-type virus or virus possessing nsp14-P203L and collected lung and nasal turbinate samples at 3 and 6 days post-infection (dpi) to analyze viral titers (Fig. 4D). We found that the nsp14-P203L virus replicated well in both the lungs and nasal turbinates of the infected hamsters at 3 and 6 dpi, and the viral titers in the lungs of the hamsters infected with the nsp14-P203L virus at 3 dpi were similar to those in the lungs of the hamsters infected with the wild-type virus (mean titers were 8.38 log unit or 8.33 log unit, respectively). We also observed clinical symptoms, including weight loss for 6 days post-infection. We found that hamsters infected with the nsp14-P203L virus showed similar weight loss to hamsters infected with the wild-type virus (Fig. 4E). The nsp14-P203L mutant virus thus grew as efficiently as the wild-type virus in hamster lungs and nasal turbinate and caused similar weight loss.

To further investigate the effect of nsp14-P203L on viral genome diversity *in vivo*, we RNA-sequenced viral genomes extracted from viruses isolated from the lungs of hamsters infected with the nsp14-P203L or wild-type virus at 6 dpi. We found that the number of mutations in the nsp14-P203L group was significantly higher than those detected in the wild-type group (63.25±12.54 mutations *versus* 32.75±3.34 mutations, respectively; *P =* 0.0066) (Fig. 4F), and that these mutations were widely distributed in the viral genomes (Fig. 4G). These results indicate that the nsp14-P203L virus replicates well in the respiratory tracts of hamsters and readily acquires more diverse mutations.

## Discussion

The mutation rate of coronaviruses, including SARS-CoV-2, is relatively low compared to other RNA viruses because coronaviruses possess an error-correcting exonuclease, nsp14. In this study, we examined mutations that occurred in nsp14, and found that SARS-CoV-2 possessing nsp14 with a proline to leucine substitution at position 203 showed a relatively higher substitution rate than SARS-CoV-2 possessing wild-type nsp14 (Tables 2 and 3; Fig. 2). Protein structure prediction analysis of SARS-CoV-2 nsp14 suggested that the P203L replacement in nsp14 decreases the binding affinity of nsp10, possibly reducing the proofreading activity (Fig. 3; Supplementary Fig. 7). Moreover, we demonstrated that a recombinant SARS-CoV-2 possessing this P203L substitution in nsp14 acquired significantly more diverse genomic mutations than the wild-type virus did during its replication in a hamster model, suggesting that mutations in nsp14, such as P203L, could accelerate the genomic diversity of SARS-CoV-2 (Figure 4).

The nsp14-P203L mutant virus grown in VeroE6/TMPRSS2 cells did not have significantly diverse genomic mutations compared to the wild-type virus, whereas the nsp14-P203L mutant virus grown in hamster lungs had significantly diverse mutations in its genome compared to the wild-type virus. One reason for this difference may be the number of genome copies of each virus in the experiments. Viruses grown in VeroE6/TMPRSS2 cells were sampled to examine genomic mutations at 2 days post-infection, whereas viruses grown in hamster lungs were assessed at 6 days post-infection. Therefore, the viruses may have replicated more in the hamsters’ lungs due to the longer incubation time. In addition, in VeroE6/TMPRSS2 cells, the nsp14-P203L mutant virus showed slightly reduced growth efficiency compared to the wild-type virus (Fig. 4A). Therefore, the wild-type virus may replicate more frequently in VeroE6/TMPRSS2 cells compared to the nsp14-P203L mutant virus, resulting accumulation of genomic mutations. In contrast, in hamster lungs, the replication efficiency of the nsp14-P203L mutant virus was similar to that of the wild-type virus (Fig. 4D). Therefore, since the genome replication frequency is almost the same between the wild-type and the nsp14-P203L mutant viruses, the reduced proofreading activity due to nsp14-P203L can be correctly detected. Overall, our findings suggest that amino acid substitutions in nsp14-P203L might affect viral properties such as genome mutation proofreading and genome replication efficiency.

Alanine substitution of an amino acid that is important for the proofreading activity of the nsp14 of SARS-CoV has been shown to increase viral genomic mutations in cultured cells (5, 11) and *in vivo* (10). In addition, Graham et al. reported that SARS-CoV is also attenuated by such a substitution (10). The effects of amino acid substitutions in nsp14 on coronavirus infections in nature have not been investigated. In our study, candidates that might affect nsp14 function were sought by focusing on amino acid substitutions in the nsp14 of epidemic viruses and comparing genomic mutation rates. However, the molecular mechanism of how amino acid mutations in nsp14 are involved in proofreading activity is not clear. In addition, the number of SARS-CoV-2 genomes, including nsp14-203L variants, was limited in our study, which limited our statistical analysis. Since the number of SARS-CoV-2 genomes deposited in the GISAID has increased (14,535,971 genomes, as of January 11, 2023), further molecular evolutionary analyses with phylogenetic correction may reveal more nsp14 mutations that affect its proofreading activity. Nevertheless, our findings demonstrate that rare amino acid substitutions (particularly leucine at amino acid 203) in coronavirus nsp14 may increase the rate of genomic mutations.

## Methods

### Coronavirus sequence data

The amino acid sequences of open reading frame 1ab (ORF1ab) of the 62 representative coronaviruses (summarized in Supplementary Table 1) were obtained from NCBI (https://www.ncbi.nlm.nih.gov/) and the GISAID EpiCoV database (https://www.gisaid.org) as reported previously (13). Aligned nucleotide sequences of SARS-CoV-2 and their metadata were obtained from the GISAID EpiCoV database as of September 7, 2020 (87,625 sequences). The coronavirus genome sequences annotated as having “Human” hosts were extracted, and sequences containing undetermined and/or mixed nucleotides were removed. The 28,082 remaining genomes were used for the analyses. The open reading frame (ORF) of each sequence was determined from the alignment sequence based on the ORF information of hCoV-19/Wuhan/Hu-1/2019 (GISAID ID: EPI_ISL_402125, GenBank ID: NC_045512.2), which was used as the reference sequence. Gap sites were removed, and the nucleotide sequences were translated. To detect amino acid substitutions for each site, the amino acid sequence after the stop codon was removed and amino acid sequences were aligned by using the L-INS-i program in MAFFT version v7.453 (26). From the sequence alignment of the nsp14 of the representative coronaviruses, the amino acids aligned at the amino acid positions of SARS-CoV-2 are shown using WebLogo 3 (27). The color of each amino acids is indicated according to its chemical properties within the default option. Note that, in the alignment sequence, gap sites were removed based on the SARS-CoV-2 nsp14 sequence.

### Molecular evolutionary phylogenetic analysis

To compute the phylogenetic tree of the nsp14 of the 62 representative coronaviruses, their nsp14 amino acid sequences were aligned by using the L-INS-i program in MAFFT version 7.453 (26). To estimate the appropriate amino acid substitution model, we used ProtTest3 version 3.4.2 (28) based on the Bayesian information criterion (BIC) values. We then computed a phylogenetic tree by using RAxML-NG version 1.0.0 (29) with rapid 1000 bootstrap replicates (30).

We also generated a phylogenetic tree of SARS-CoV-2 genomes belonging to the PANGO lineage B.1.1.33 including nsp14-P203L variants. Using CD-HIT-EST version 4.8.1 (31), identical genomic sequences were removed: from 200 to 142 sequences. The 142 nucleotide sequences were aligned by using the L-INS-i program in MAFFT version 7.453 (26). To estimate the appropriate nucleotide mutation model, we used ModelTest-NG (32) based on the Akaike information criterion (AIC) values, and GTR+I+G4 was selected. We then computed a phylogenetic tree by using RAxML-NG version 1.0.0 (29) with rapid 1000 bootstrap replicates (30).

### Substitution rate analysis

A multiple sequence alignment of SARS-CoV-2 genomes obtained from the GISAID database was used to analyze the genomic evolutionary rate. The number of nucleotide mutations per site from between sequences was counted. Analyses were conducted using the Kimura 2-parameter model. Codon positions included 1st+2nd+3rd+Noncoding. All gaps were removed for each sequence pair with a pairwise deletion option. The number of nucleotide mutations was counted in MEGA X (33). The hCoV-19/Wuhan/Hu-1/2019 (GISAID ID: EPI_ISL_402125, GenBank ID: NC_045512.2) was selected as the reference sequence. Sequences with information that included the collection date were used for the analysis. Note that we omitted sequences that were collected before December 26, 2019, the date on which the reference sequences were collected. The approximate straight line was calculated by using the least-squares method and in-house python scripts.

### Measurement of natural selection pressure

TranslatorX (34) was used to align the nucleotide sequences based on the aligned amino acid sequences of the nsp14 of the representative coronaviruses. To determine the evolutionary selection pressures on the nsp14 gene in coronaviruses, we computed the ratio of nonsynonymous substitution rates (dN) and synonymous substitution rates (dS) per site (dN/dS) by using Datamonkey software (15, 35) through MEGA X (33).

### Cells

VeroE6/TMPRSS2 (JCRB 1819) cells were maintained in Dulbecco’s Modified Eagle Medium (DMEM) supplemented with 10% FBS, 1% Penicillin-Streptomycin Solution, and 1 mg/ml geneticin (G418; Nacalai Tesque, Cat# 09380-44). HEK293-hACE2/hTMPRSS2 cells were maintained in DMEM supplemented with 10% FBS and 1% Penicillin-Streptomycin Solution. VeroE6/TMPRSS2 and HEK293-hACE2/hTMPRSS2 cells were maintained at 37℃ with 5% CO_2_.

### Reverse genetics

Recombinant SARS-CoV-2 was generated by using circular polymerase extension reaction (CPER) as previously described (25). Briefly, nine DNA fragments encoding the partial genome of SARS-CoV-2 (hCoV-19/Japan/TY-WK-521/2020) were amplified by PCR using PrimeSTAR GXL DNA polymerase (Takara, Cat# R050A). A linker fragment encoding the hepatitis delta virus ribozyme, the bovine growth hormone poly A signal, and the cytomegalovirus promoter was also amplified by PCR. The ten DNA fragments were mixed and used for CPER (25). To produce recombinant SARS-CoV-2 (seed virus), the CPER products were transfected into HEK293-hACE2/hTMPRSS2 cells by using TransIT-LT1 (Takara, Cat# MIR2305). At one day post-transfection, the culture medium was replaced with DMEM (high glucose) (Nacalai Tesque, Cat# 08459-64) containing 2% FBS and 1% Penicillin-Streptomycin Solution. At 4–10 days post-transfection, the culture medium was collected and stock at -80℃. The viruses were used to inoculate VeroE6/TMPRSS2 cells to produce stock viruses. No mutations were found in the wild-type viruses, whereas the nsp14-P203L viruses were found to contain mixed unexpected mutations such as C12119T/C (nsp8-P10S/P) and C17143T/C (nsp13-R303C/R). To obtain viruses with the single mutation of nsp14-P203L, a plaque-purified virus clone was grown in VeroE6/TMPRSS2 cells. All experiments with transfectants generated by reverse genetics were performed in an enhanced biosafety level 3 (BSL3) containment laboratory approved for such used by Osaka university.

### Deep sequencing analysis

Viral RNA was extracted from samples by using the QIAamp Viral RNA Mini kit (Qiagen) according to the manufacturer’s instructions. The extracted RNA was reverse transcribed with ProtoScript II (NEB) using N6 primers. The cDNA was converted to double-stranded DNA by NEBNext Second Strand Synthesis (NEB). In addition, double-stranded DNA was fragmented by the SureSelect Fragmentation Enzyme. Fragmented DNA was made into an NGS library with the SureSelect XT Low Input Kit. In addition, hybridization was performed using a SARS-CoV-2 custom probe, and sequence-ready libraries were sequenced in a paired-end run using by MiSeq (Illumina).

The raw FASTQ format data files that were obtained from the sequencing samples were filtered using cutadapt version 3.2 (36) to remove low-quality reads and adapters. Then, the filtered sequences were mapped into the reference SARS-CoV-2 genome (Wuhan-Hu-1, NC_045512.2) by using BWA version 0.7,17-r1188(37). Variant calling was done using MuTect2 in GATK suite version 4.2 (38), and nucleotide changes were counted by using in-house python scripts.

### Animal experiments and approvals

All animal experiments were approved by the Animal Research Committee of the Research Institute for Microbial Diseases, Osaka University (assurance number, R03-04-0). All animals were housed under specific pathogen-free conditions in a temperature and humidity control environment with a 12 h:12 h light dark cycle and ad libitum access to water and standard laboratory chow. Virus inoculations were performed under anesthesia, and all efforts were made to minimize animal suffering.

These *in vivo* studies were not blinded, and animals were randomly assigned to infection groups. No sample-size calculations were performed to power each study. Instead, sample size was determined based on prior *in vivo* virus challenge experiments.

### Experimental infection of Syrian hamsters

Seven-week-old male wild-type Syrian hamsters (Japan SLC Inc., Shizuoka, Japan) were used in this study. Baseline body weights were measured before infection. Under isoflurane anesthesia, four hamsters per group were intranasally inoculated with 10^3^ PFU in 30 μl of recombinant virus. Body weight was monitored daily for 6 days after infection. For virological examinations, four hamster per group were intranasally infected with recombinant viruses; 3 and 6 dpi, the animals were euthanized, and nasal turbinate and lungs were collected. The virus titers in the nasal turbinate, trachea and lungs were determined by performing plaque assays on VeroE6/TMPRSS2 cells.

### *In silico* structural analysis

A PDB file (7EGQ) of the SARS-CoV-2 replication-transcription complex (RTC) was obtained from the PDB database (https://www.rcsb.org). Using the nsp14-nsp10 complex (chains H and K) of the structure, further analyses were performed. The residues at the interface of the SARS-CoV-2 nsp10-nsp14 docked complex were determined using the “InterfaceResidues” PyMol script (http://www.pymolwiki.org/index.php/InterfaceResidues/). We used the SARS-CoV-2 nsp10-nsp14 docked complex to predict the effect of the P203L mutation on protein-protein interactions by using the BeAtMuSiC program version 1.0 (http://babylone.3bio.ulb.ac.be/beatmusic/query.php) (39). P203L mutant generation, hydrophobicity calculation, and visualization were performed using PyMol version 2.4.0.

### Statistical analysis

All data were analyzed in GraphPad Prism software (version 9.4.0). Statistical analysis included ANOVA with multiple corrections, post-tests, or an unpaired Student’s t test. Differences among groups were considered significant for *P* values < 0.05.

## Acknowledgments

We thank Dr. Susan Watson for scientific editing. We acknowledge the NGS Core laboratory, Genome Information Research Center, Institute for Microbial Diseases, Osaka University, for assistance with the RNA sequence. We thank the GISAID EpiCoV database (https://www.gisaid.org), which shares sequence data of SARS-CoV-2 and related viral species, and all researchers who deposited SARS-CoV-2 genomes and related information in GISAID (https://epicov.org/epi3/epi_set/230110sz). This study was supported by KAKENHI Grants-in-Aid for Scientific Research on Innovative Areas 16H06429 (to Y.M., T.W., and S.N.), 16K21723 (to Y.M., T.W., and S.N.), JP16K21723 (to Y.M.), JP16H06432 (to Y.M.), 16H06434 (to T.W), and 19H04843 (to S.N.); Grants-in-Aid for Scientific Research (B) JP22H02521 (to T.W.) and JP21H02736 (to C.O); Grant-in-Aid for Early-Career Scientists 22K15469 (to K.T.); Grant-in-Aid for JSPS Fellows 21J01036 (to K.T.); AMED Research Program on Emerging and Re-emerging Infectious Diseases JP20fk0108281 (to Y.M.), JP19fk0108113 (to Y.M. and T.W.), JP20fk0108401 (to C.O.), JP21fk0108493 (to C.O.), and 19fk0108171 (to S.N.); AMED Cyclic Innovation for Clinical Empowerment (CiCLE) JP20pc0101047 (to Y. M.); AMED under Grant Number JP223fa627002h (to T.W.); AMED Advanced Research and Development Programs for Medical Innovation (AMED-CREST) 22gm1610010h0001 (to T.W.); 2020 Tokai University School of Medicine Research Aid (to S.N.); Japan Science and Technology Agency (JST) CREST JPMJCR20H6 (to S.N.); JST Moonshot R&D JPMJMS2025 (to Y.M.); the Tokyo Biochemical Research Foundation (to T.W.); the Takeda Science Foundation (to T.W.); and RIKAKEN HOLDINGS CO. Young Researcher Support Grant-in-aid (to K.T.). The super-computing resources were provided by the Human Genome Center (the Univ. of Tokyo) and the NIG supercomputer at ROIS National Institute of Genetics.

## Contributions

K.T., T.W. and S.N. conceived the study. K.T., M.T.U and S.N analyzed data. C.O. and Y.M. established 293-hACE2/hTMPRSS2 cells. K.T. generated recombinant viruses. K.T. and Y.K. performed in vitro experiments. K.T., S.S. and T.W. performed animal experiments. K.T, M.T.U., T.W and S.N. wrote the manuscript. All authors read and approved the final manuscript.

## Competing interests

The authors have no competing interests.

## Data Availability Statements

The data underlying this article will be shared upon reasonable request to the corresponding author. All RNA-seq data sequenced in this study were deposited in DDBJ Sequence Read Archive (DRA) with the following accession numbers: DRR411740 - DRR411753

## Notes

### Competing Interest Statement

The authors have declared no competing interest.

### Summary of Updates

Figure 2 was replaced with Figure S4; Figure 3 was revised using SARS-CoV-2 nsp14; some discussion including limitation of this study was added.

