## Supplementary Figures for "Genomic diversity of SARS-CoV-2 can be accelerated by mutations in the nsp14 gene"

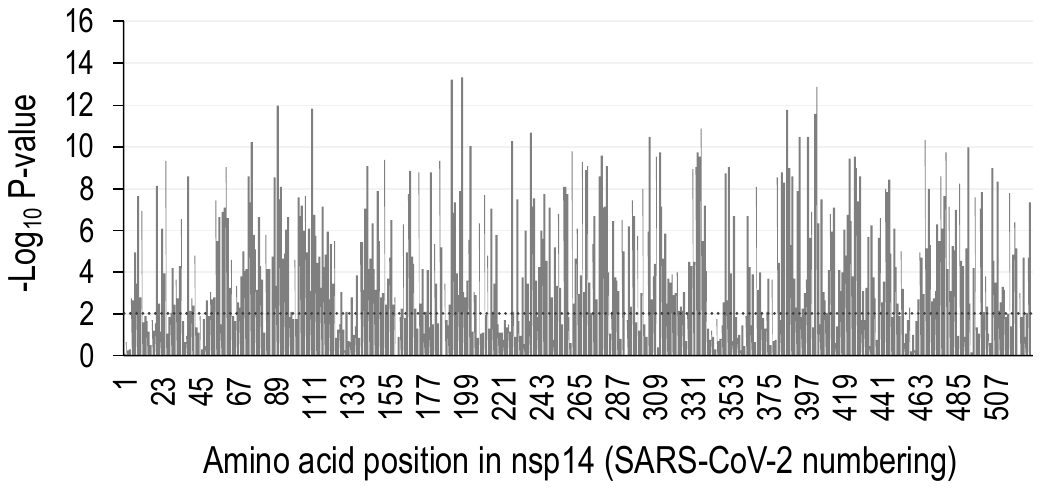

**Supplementary Fig. 1: Negative selection profiles of nsp14 encoded by the 62 representative coronavirus genomes**

The abscissa indicates the codon positions, with the scale bar at the top. The ordinate indicates the (–log_10_ P) value for each position when dN/dS < 1. Each position of nsp14 is shown according to the numbering of SARS-CoV-2. The codon position 504 of nsp14 was calculated by using 61 CoVs excluding the Chinese waterside skink CoV. Dotted line indicates *P* = 0.01.

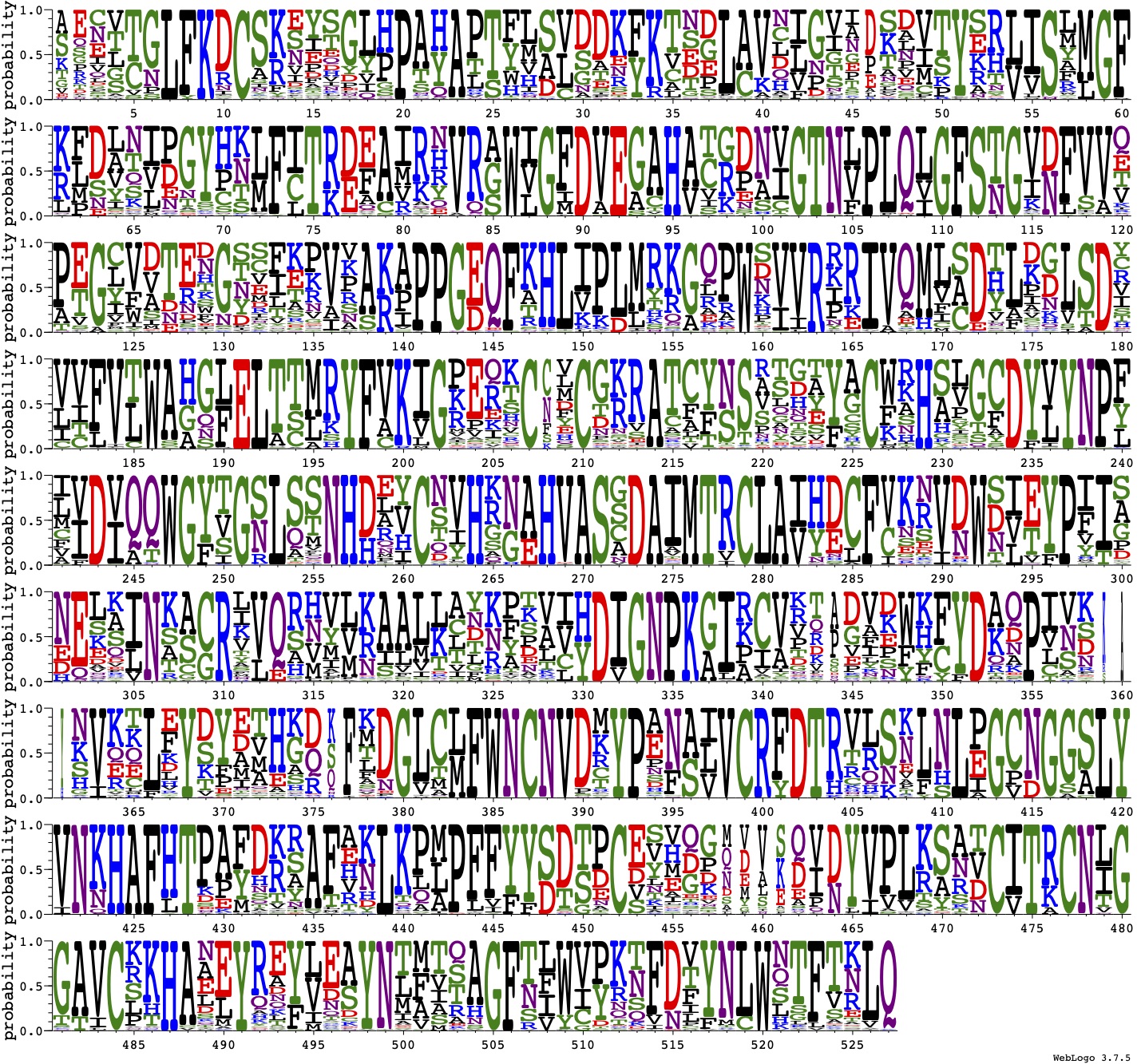

**Supplementary Fig. 2: WebLogo highlighting conserved amino acids in the nsp14 of 62 representative coronaviruses**

From the sequence alignment of the nsp14 of the 62 representative coronaviruses, the amino acids aligned at the amino acid positions of SARS-CoV-2 are shown.
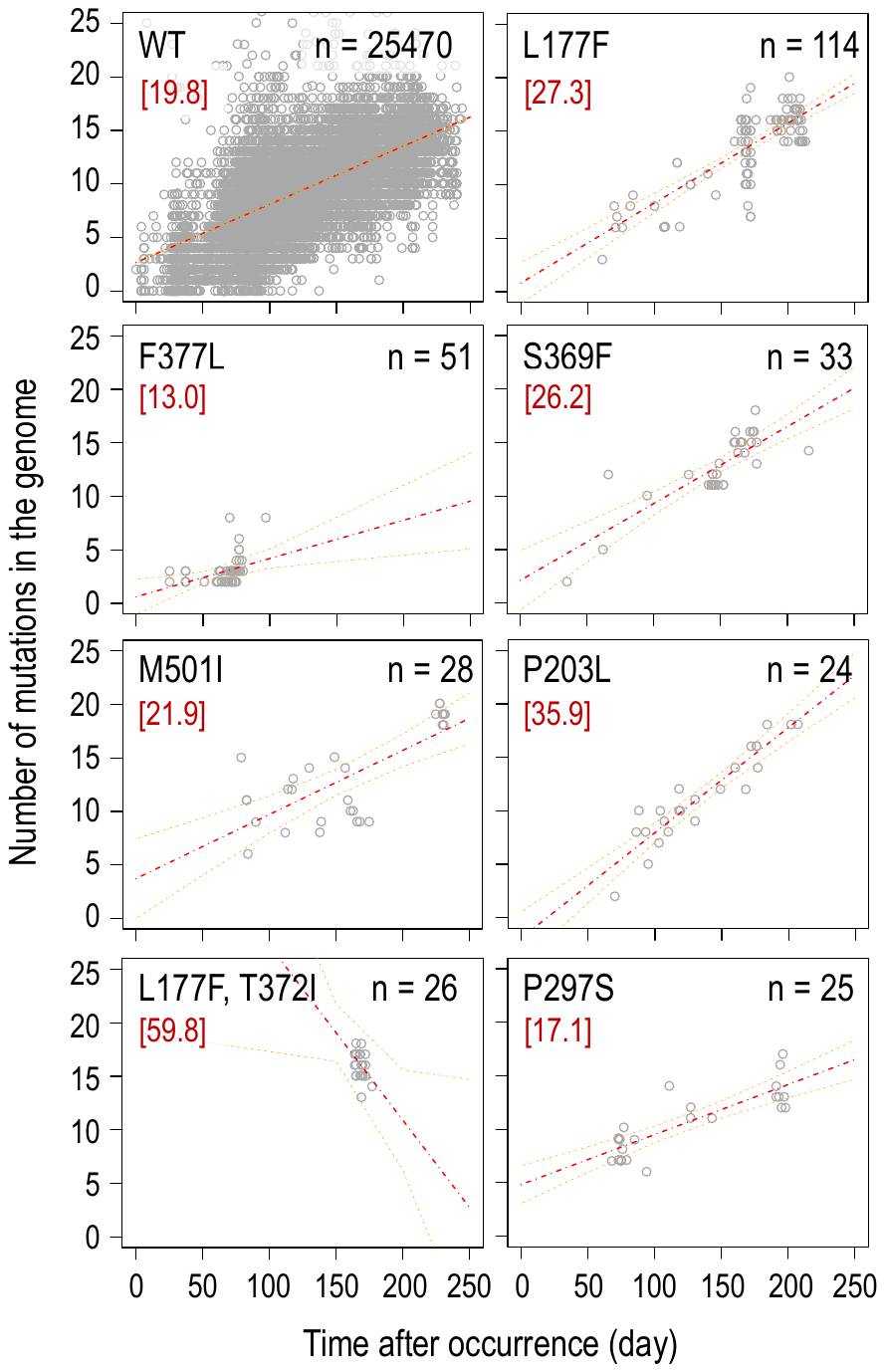

**Supplementary Fig. 3: Scatter plot of nucleotide mutations in SARS-CoV-2 genomes**

Genome diversity of SARS-CoV-2 containing wild-type (WT) nsp14 and 7 nsp14 mutants is shown. The X-axis indicates the sampling date and the Y-axis indicates the number of nucleotide mutations in the genomes. Mutation rates per year in the genome are shown in red letters. The red and orange dotted lines correspond to the regression line and 95 % confidence intervals, respectively.

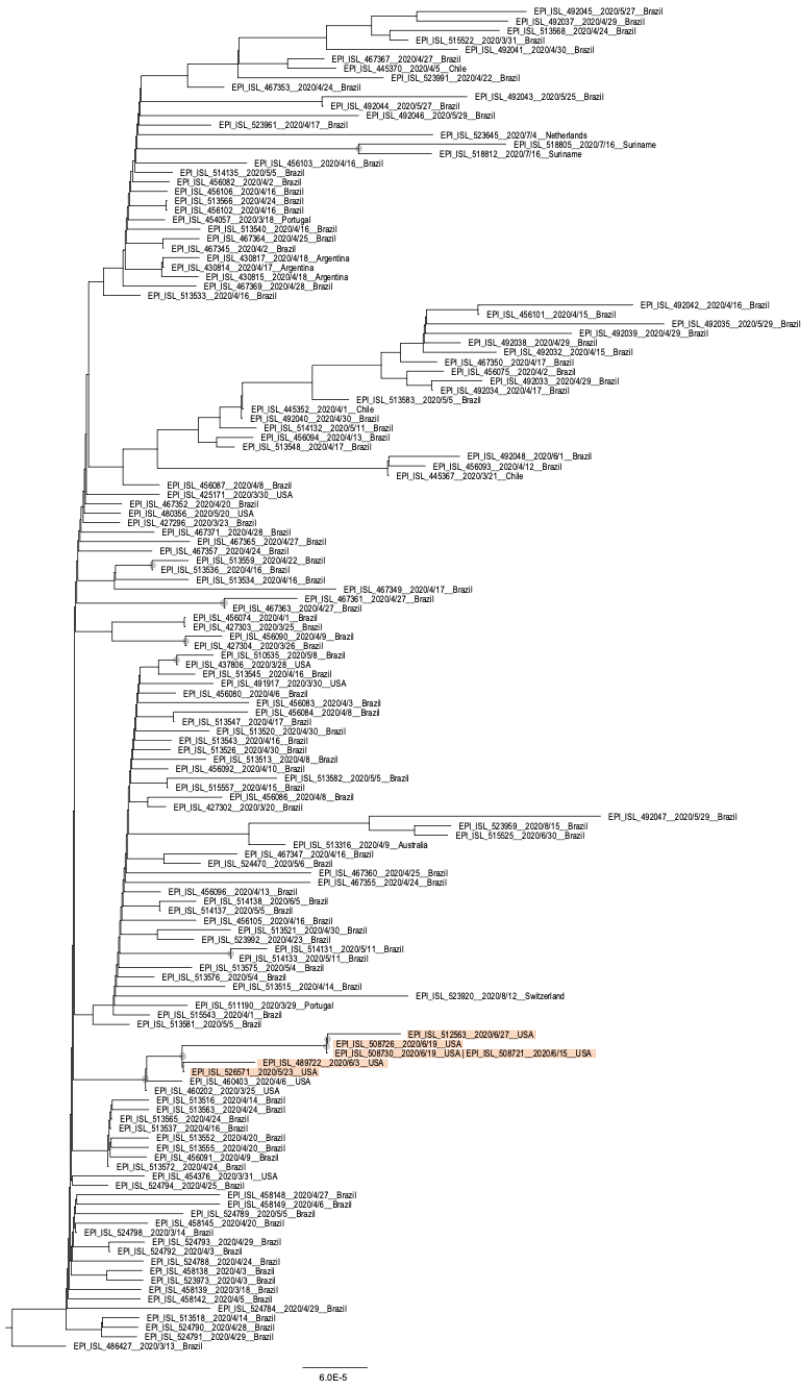

**Supplementary Fig. 4: Phylogeny of nsp14-P203L variants in the PANGO lineage B.1.1.33**

ML tree of SARS-CoV-2 genomes in the B.1.1.33-lineage. The nsp14-P203L variant is highlighted in light red. A gray circle corresponds to bootstrap values ≥ 70%.

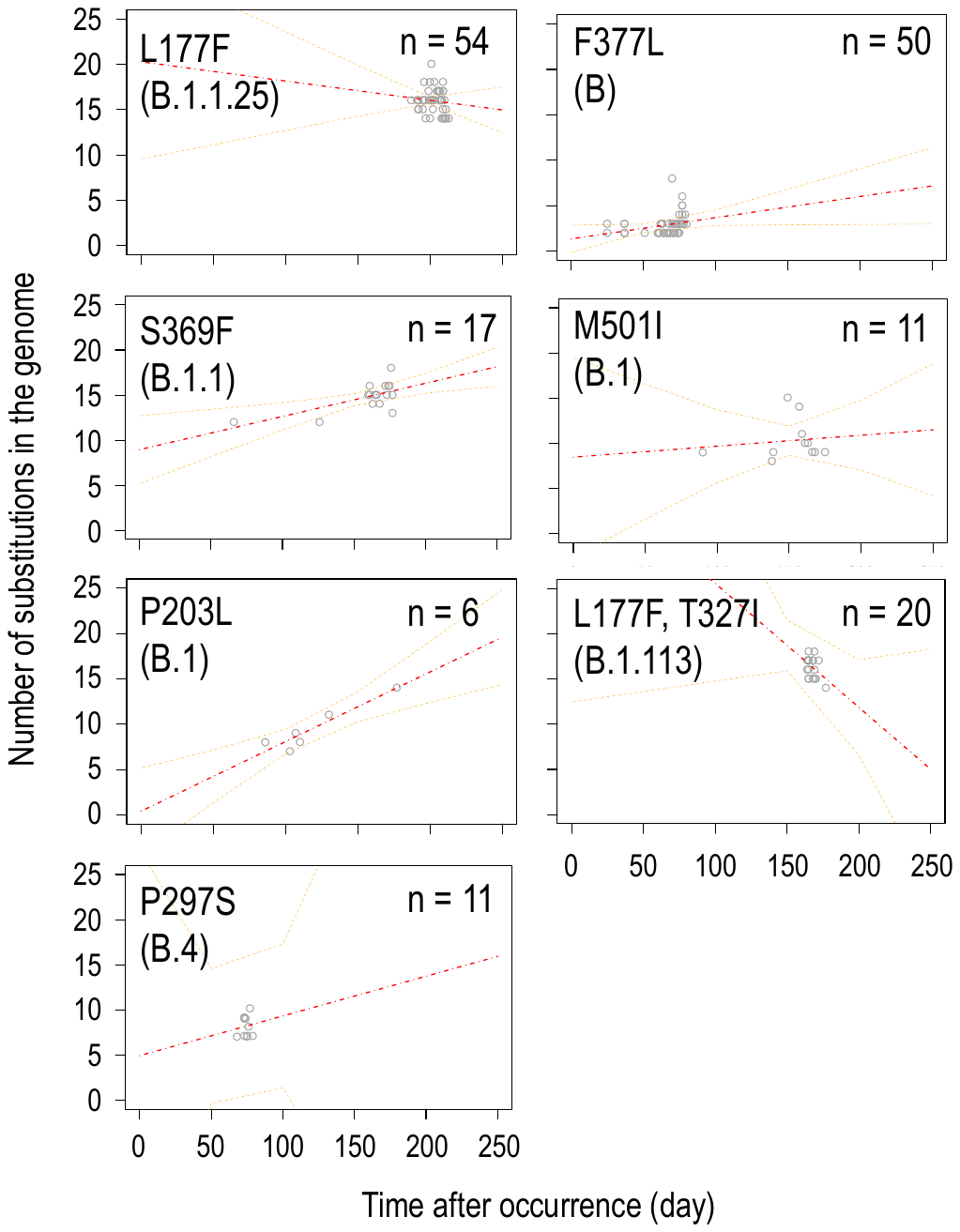

**Supplementary Fig. 5: Scatter plot of the 7 nsp14 variants representing each cluster where the maximum number was observed**

Nucleotide mutation rates of 7 nsp14 variants for each cluster. The X-axis indicates the sampling date and the Y-axis indicates the number of nucleotide mutations in the genomes. The red and orange dotted lines correspond to the regression line and 95 % confidence intervals, respectively.

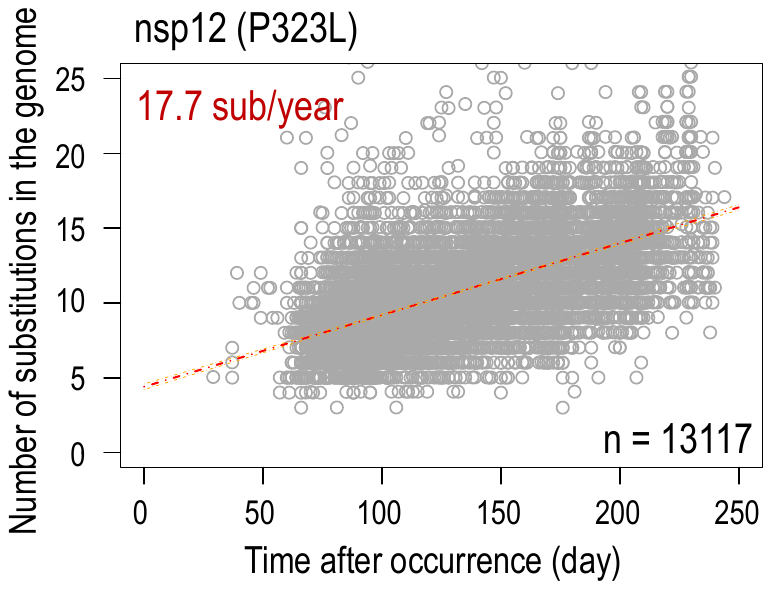

**Supplementary Fig. 6: Scatter plot of nucleotide mutations in nsp12-P323L variants**

Genome diversity of nsp12-P233L variants is shown. Duplicate sequences were removed, leaving only the sequences with the oldest sampling date. The X-axis indicates the sampling date and the Y-axis indicates the number of nucleotide mutations in the genomes. Mutation rates per year in the genome are shown in red letters. The red and orange dotted lines correspond to the regression line and 95 % confidence intervals, respectively.

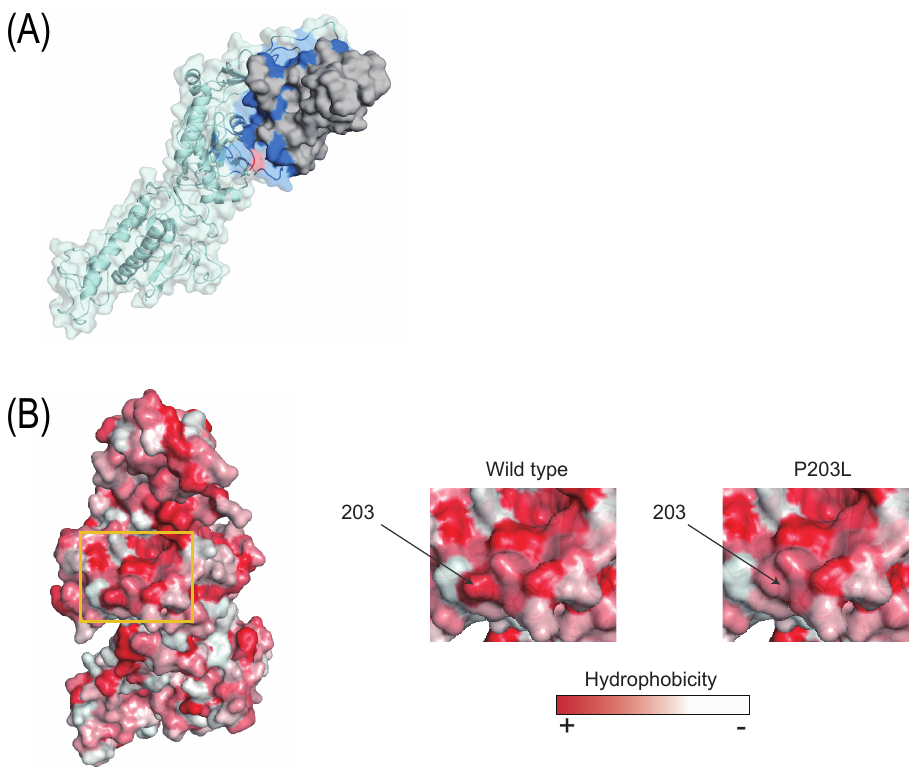

**Supplementary Fig. 7: Three-dimensional position of the P203L mutation in nps14 and the interaction site with nsp10**

**(**A) Amino acid position on the three-dimensional structure of nsp14. The nsp14 and nsp10 is shown in light blue and gray, respectively. The site of interaction with nsp10 is shown in dark blue. Amino acid position 203 in nsp14 is shown in red. (B) Hydrophobic (red) and hydrophilic regions (white) at the surface of the nsp14-203P (wild type) or nsp14-203L protein.
